# Cryo-EM structures of an *Acinetobacter baumannii* podovirus reveal a canonical capsid maturation

**DOI:** 10.64898/2026.11.06.765006

**Authors:** Heng Xue, Guibo Rao, Xinfeng Li, Mingyue Zhong, Sheng Cao, Hang Yang

## Abstract

Phages that hunting multidrug resistant bacteria represent a promising approach in precision medicine nowadays, including these infecting *Acinetobacter baumannii*. To data, a large number of *A. baumannii* phages has been isolated, however, complete structural information remains not adequately released. Previously, we reported a podovirus, AbP20, that infects *A. baumannii* with potent efficacy in mouse model. In the present work, we further report the high-resolution cryo-electron microscopy (cryo-EM) structures of AbP20 and its procapsid. The near-atomic resolution reveals that AbP20 is composed of five distinct structural subunits, i.e., capsid, portal, nozzle, fiber, and adaptor. We then characterized the assembly mechanisms of molecular interactions within capsid and portal-tail complex. By co-expressing the scaffold protein and the capsid protein of AbP20, we constructed the self-assembled DNA-free procapsid of AbP20. A near-atomic resolution reconstruction reveals it a double-layered concentric spherical architecture with a nearly 1:1 ratio of these two proteins. Comparison between the mature and apo-form of the capsid of AbP20 uncovers a canonical model of conformational switch from procapsid to capsid. Altogether, the present work reports the first whole atom structure of a podovirus infecting *A. baumannii*, providing a reference for structure-based phage engineering and editing.

**Author summary:** Multidrug-resistant bacteria, particularly *Acinetobacter baumannii*, pose a severe threat to global health. As a promising alternative in precision medicine, phages, viruses that naturally infect and kill bacteria, offer a new avenue to combat these dangerous infections. Although numerous phages targeting *A. baumannii* have been identified, their detailed structural information remains largely elusive. In this study, we utilized high-resolution cryo-electron microscopy (cryo-EM) to determine the native virion structure of AbP20, a podovirus with potent efficacy against *A. baumannii*. We revealed the detailed architecture and assembly of its distinct components. Furthermore, by reconstructing the procapsid structure of AbP20, we elucidated the role of its scaffold protein in capsid assembly and maturation. This work reports the first complete atomic structure of a podovirus infecting *A. baumannii*, reveals a canonical model for virion capsid maturation, and provides a valuable reference for structure-guided phage engineering and editing.

## Introduction

The frequent emergence and transmission of antimicrobial resistant pathogens pose critical threats to global health. Multidrug-resistant, even pan-drug-resistant *Acinetobacter baumannii* has achieved worldwide prevalence, with a particularly pronounced impact on patients with ventilator-associated pneumonia and bloodstream infections [1], rendering it one of the foremost challenges in the global fight against infectious diseases. Phage therapy has been revisited as a key strategy to address this challenge. Currently, clinical phage therapy has been successfully employed in the treatment of multiple cases of *A. baumannii* infection [2–5], yielding encouraging outcomes.

To date, a large number of *A. baumannii* phages have been isolated and characterized. Most of them are caudovirus with an icosahedral or elongated icosahedral capsid. However, only a very limited number of *A. baumannii* phage structures have been resolved, for instance, the non-tailed phage AP205 [6], and the siphovirus Mystique [7] and SH-Ab15497 (PDB: 9K4G, 8XB6). Structural information on the podovirus and myovirus that infection *A. baumannii* is still largely inadequately addressed. The scarcity of such information impedes a deeper understanding of their virion assembly process, especially the dramatic conformational changes and structural rearrangements occurred in capsid maturation.

It is believed that virion capsid assembly begins with the formation of a DNA-free precursor, i.e., the procapsid that consists of an inner scaffold core and an outer capsid shell, for many tailed phages, such as *Salmonella* phage P22, *B. subtilis* phage SPP1, *E. coli* phage T4, T7, and λ [8–10]. More specifically, the assembly of icosahedral capsid in tailed phages depends on a transient internal scaffold element, which acts as chaperone-like factor and is absent from the mature virion. These scaffold elements can be either independent proteins, i.e., scaffold protein as observed in phage P22, SPP1, and T7, or N-terminal fusion domains of the major capsid protein (MCP) as observed in *E. coli* phage T4 and λ [11, 12]. The removal of scaffold protein results in MCP assembling and the finalized capsid maturation [13–15], within which the interactions of scaffold protein with capsid or portal protein make a critical role. In certain phages, such as P22 and T7, the exit of scaffold protein is accompanied by the ATP-driven

DNA packaging process, afterwards, the icosahedral shell of the procapsid undergoes dramatic conformational changes and matures into the larger, more angular shell typical of infectious phages [16–18]. However, the transition from the procapsid to the mature capsid and the key role of scaffold protein for *A. baumannii* phages remain to be further elucidated.

Previously, we reported a lytic podovirus, AbP20, infecting *A. baumannii* shows good therapeutic potential [19]. Here, we report the structures of the native virion and procapsid of AbP20 using single-particle cryo-electron microscopy (cryo-EM), with detailed characterization of the architecture of its head and tail component, providing new insights into understanding the assembly mode and maturation mechanism of *A. baumannii* phages.

## Results

### Overview of the whole AbP20 virion

AbP20 phage was propagated in *A. baumannii* strain Ab28, and high purity phage particles (~10^13^ PFU/mL) were obtained by density gradient ultracentrifugation (S1 Fig). Using cryo-EM single-particle analysis, we determined high-resolution structures of the AbP20 capsid and the portal-tail complex at global resolution of 3.0 Å and 3.2 Å, respectively (S2 Fig). The AbP20 virion exhibits an icosahedral symmetry-like capsid with a diameter of approximately 560 Å and a short, non-contractile tail measuring ~160 Å in length. (Fig 1A). The overall structure of AbP20 comprises five distinct protein subunits (Fig 1B). Specifically, the entire AbP20 capsid is composed of 415 copies of the capsid protein (gp37), which assemble into 60 hexons and 11 pentons (Fig 1B). The capsid shell encloses a large internal cavity that accommodates densely packed viral DNA (Fig 1A). The tail apparatus consists of a dodecameric portal protein (gp35), a dodecameric adaptor protein (gp39), a hexameric nozzle protein (gp40), and a trimeric fiber complex (gp44) arranged in a hexameric pattern (Fig 1B-C). A canonical 12-copy structure portal protein is anchored to the inner surface at one pentameric vertex of the capsid and extends outward to interact with the adaptor protein (Fig 1A). The intra-capsid portion of the portal is surrounded by concentric layers of viral DNA (Fig 1A), a structural feature commonly observed in tailed phages [20, 21]. Together with the ring-shaped adaptor and the tubular nozzle, the external end of the portal forms a continuous channel for DNA ejection (Fig 1B). Notably, the distal portion of the tail fiber could not be fully modeled due to intrinsic conformational flexibility, which precluded the acquisition of high-resolution atomic structure.

**Fig 1.**
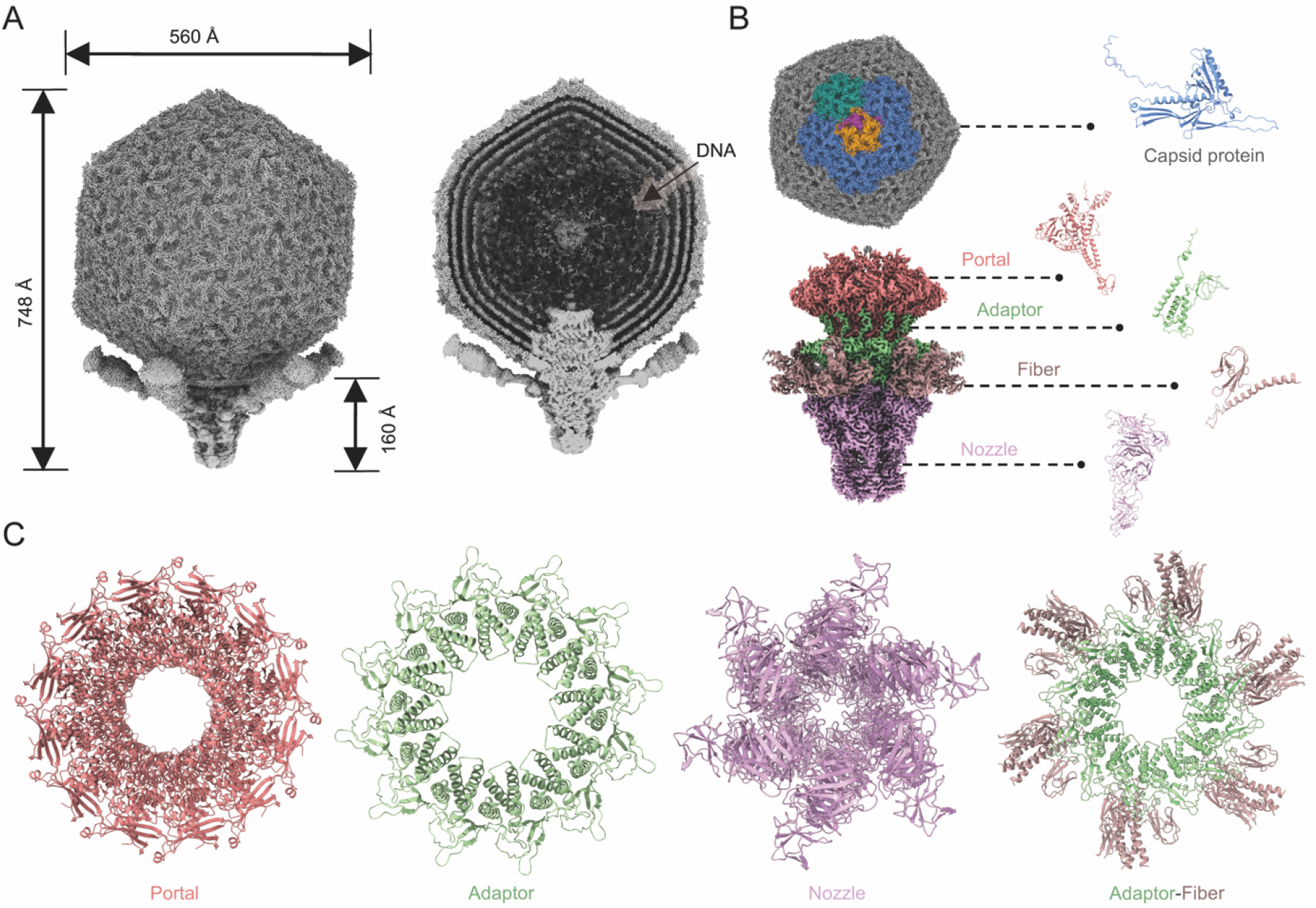
The overall architecture of the AbP20 virion. (**A**) Surface (left) and section (right) views of the AbP20 virion. Packed DNA inside the capsid is indicated by an arrow. (**B**) Details of the structural components and monomer structures of the structural proteins, including the major capsid protein (blue), portal (blush), adaptor (green), fiber (brown), and nozzle (purple).(**C**) Top views of the symmetry rings formed by various phage proteins, including the portal, adaptor, nozzle, and adaptor-fiber complex.

### The capsid structure of AbP20

Each vertex of the AbP20 icosahedral capsid features a central penton surrounded by five hexons, forming five asymmetric units (Fig 2A). Each asymmetric unit of the AbP20 capsid consists of one hexameric and one pentameric subunit, with a triangulation number (T) of 7.

**Fig 2.**
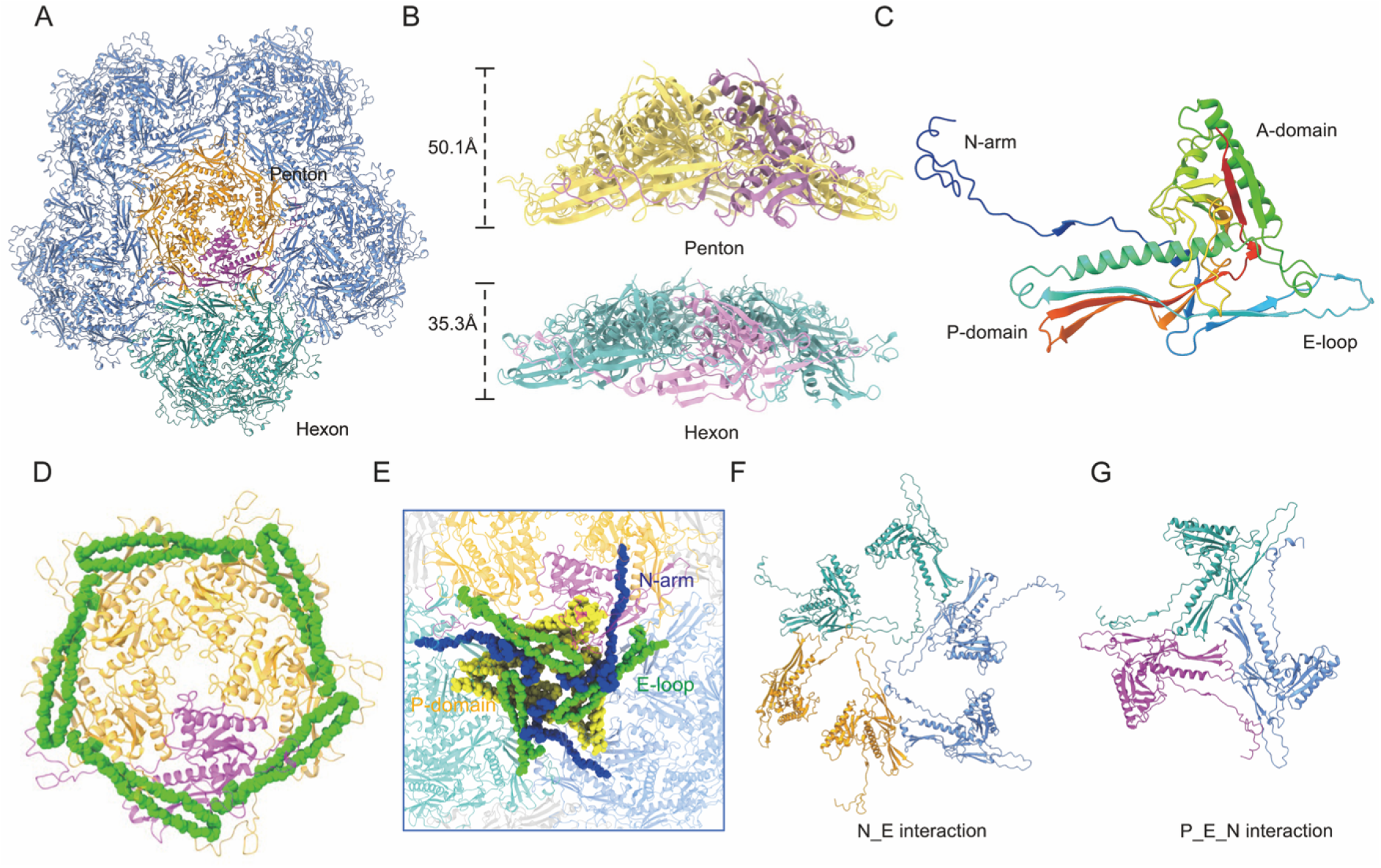
Overall structure of AbP20 capsid. (**A**) Each vertex of the AbP20 capsid comprises one central penton (orange) and five surrounding hexons (green). An asymmetric unit (T = 7) consisting of one hexon (green) and one penton subunit (orange) is highlighted. (**B**) Side views of the penton and hexon, with heights of 50.1 Å and 35.3 Å, respectively. (**C**) Structure of the capsid protein gp37, with domains colored differently: N-arm (blue), A-domain (green), P-domain (orange), and E-loop (cyan). (**D**) The A-domains form the structural core of the capsid, while the E-loops wrap around adjacent N-arms and P-domains, creating a circular interaction network that enhances capsomer stability. (**E**) Interactions between adjacent E-loops and P-domains, further reinforced by extended N-arms engaging multiple subunits, secure neighboring structures. (**F**) N-arm-E-loop interactions between an adjacent penton and two hexons. (**G**) P-domain-E-loop-N-arm interactions between an adjacent penton and two hexons.

The AbP20 capsid achieves precise radial thickness regulation (50.1 Å and 35.3 Å for pentons and hexons, respectively) through differential domain orientation and extension of gp37 at the penton and hexon positions (Fig 2B). The capsid protein gp37 comprises a long N-terminal arm (N-arm), an extended loop (E-loop), a peripheral domain (P-domain), and an axial domain (A-domain) (Fig 2C), adopting a typical folding that was first identified in phage HK97 [22, 23]. Since such folding pattern has also been observed in numerous tailed phages, including T7, as well as in herpesviruses [16, 24], we then carried out a pairwise sequence alignments between the AbP20 capsid protein and those of HK97 (PDB: 1OHG), T7 (PDB: 3J7W), and herpesvirus (PDB: 6CGR) and found a high degree of conservation in tertiary architecture, although they share a relatively low overall sequence similarity (S3 Fig).

Next, we described the interactions *via* which the capsid maintain stability. It turned out that both pentons and hexons are stabilized by similar intra-capsomeric interactions, specifically, the A-domain serves as the central scaffold, while the E-loop adopts a clasping conformation, wrapping around the adjacent N-arm and P-domain, forming a closed ring like structure (Fig 2D and S4A Fig). The inter-capsomeric interactions are mediated by mutual penetration of the E-loops and P-domains of neighboring capsomers and further reinforced by the extended N-arms interacting with multiple subunits (Fig 2E and S4B Fig). At the inter-capsomeric level, we identified two distinct types of interfaces. The first is called the N_E interaction, i.e., the N-arm inserts into the groove of the adjacent E-loop, serving as a primary docking module (Fig 2F); the second is called the P_E_N interaction, representing a tripartite interlocking hub formed by the P-domain, E-loop, and N-arm from adjacent subunits (Fig 2G). This arrangement results in mutual penetration of the E-loops and P-domains of neighboring capsomers, while the extended N-arms bridge across the interface to bind multiple subunits from the opposing capsomer (S4C-D Fig).

### The portal-tail complex of AbP20

The AbP20 portal-tail complex replaces one penton of the icosahedral capsid. The portal protein gp35 is embedded within the capsid, stabilized by the adaptor protein gp39, and capped distally by the nozzle protein gp40 (Fig 3A). The portal-tail complex possesses a central channel with a gradually decreasing inner diameter and a negatively charged inner surface (S5A-B Fig). The portal protein gp35 exhibits an alternating arrangement of positive and negative charges, which may serve to stabilize the genomic DNA inside the capsid [25]. The nozzle region carries a strong negative charge to facilitate DNA injection (S5B Fig). Each portal subunit is mainly composed of four canonical domains: crown, wing, stem, and clip (S5C Fig), in which the β-hairpin in the wing domain connects the corresponding β-hairpin of the neighboring subunit in a head-to-tail manner, forming a structure that wraps the portal dodecamer.

**Fig 3.**
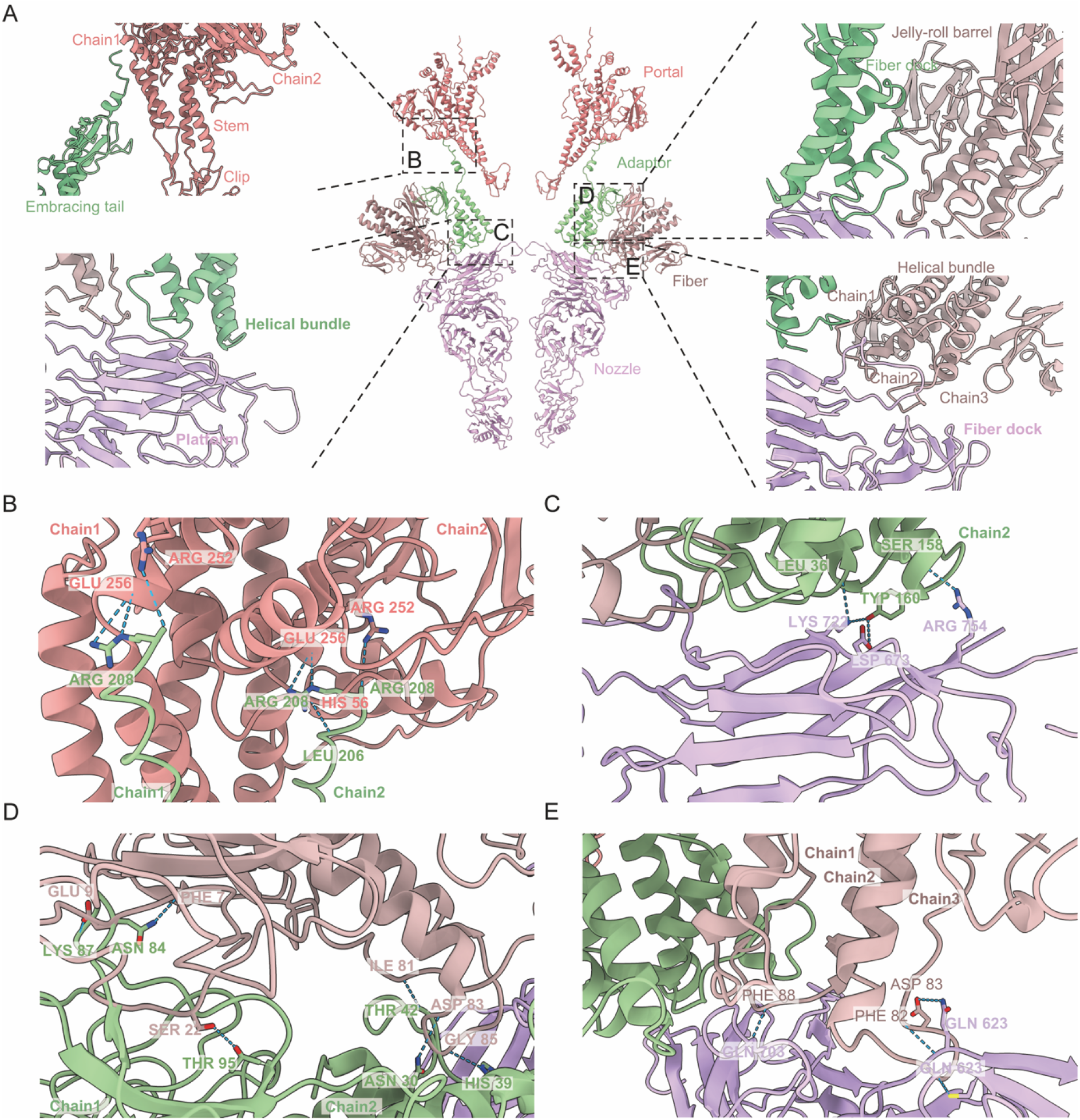
Structure of the portal-tail complex of AbP20. (**A**) Enlarged views of the portal–adaptor, adaptor–nozzle, and adaptor–nozzle–fiber interfaces, with the relevant interfaces highlighted by dashed boxes. The regions involved in the interactions are labeled. (**B-E**) High-magnification views showing the interaction details of the four interfaces. Residues involved in the interactions are shown as sticks and labeled. Hydrogen bonds are indicated by blue dashed lines.

Each subunit of the adaptor comprises a helical bundle, an embracing tail domain, and a capsid-docking domain (S5C Fig), exhibiting a domain arrangement similar to that of the adaptor of T7 phage [26]. The C-terminal tail extends into the cleft between two portal protein gp35 subunits for connection (Fig 3A). Each subunit of the nozzle is featured with a central β-propeller domain composed of seven blades, which bridges a platform domain and a fiber dock domain at one side to a tip domain at the other side (S5C Fig). The tip region could directly interact with the host outer membrane during phage infection to mediate DNA injection [27]. At the junction between the adaptor and the nozzle, six tail fibers (trimeric gp44) are symmetrically attached (Fig 3A). Because the distal portion of the fiber is flexible and not visible in the cryo-EM density map, we could only resolve the structural information for its N-terminal residues 1-126.

The interwoven protein chains generate complex interactions among the portal, adaptor, nozzle, and fiber (Fig 3B-E). We performed interface analysis among them. We performed interface analysis on these components. Each pair of portals forms an interaction interface with two adaptors, comprising 7 hydrogen bonds (Fig 3B). Four hydrogen bonds could be observed between the adaptor and nozzle (Fig 3C). The N-terminus of the trimeric tail fiber interacts with two adaptors through a jelly-roll domain, forming 7 hydrogen bonds distributed across two spatially distant interaction interfaces (Fig 3D), and the nozzle via a helical bundle, forming an interaction interface with 4 hydrogen bonds (Fig 3E).

### Structure of the procapsid discloses a conformational transition

To elucidate the mechanism of AbP20 capsid maturation, we co-expressed the AbP20 capsid protein (gp37) and scaffold protein (gp36) in a prokaryotic system, yielding self-assembled procapsids (S6A Fig), which consistent with the morphology of naturally occurring procapsids obtained from the phage lysates (S6B Fig). Cryo-electron microscopy (cryo-EM) combined with icosahedral reconstruction enabled us to determine the icosahedral structure of the procapsid particles at a resolution of 3.2 Å (Fig 4A and S7 Fig). Structural analysis uncovered a double-layered concentric spherical architecture within the procapsid (Fig 4B and S8A Fig), which likely accommodates 1:1 ratio of the capsid protein gp37 to the scaffold protein gp36. Moreover, the U-shaped helix-turn-helix motif of the scaffold protein was clearly observed inside the AbP20 procapsid; whilst the rest of the internal densities did not support high-resolution reconstruction. We modeled the C-terminal residues (242-286) of the scaffolding protein gp36 into the U-shaped structure (Fig 4C-D). Two C-terminal α-helices interact with the N-arms of two gp37 subunits, thereby stabilizing the procapsid structure. Within the scaffold region, H259 forms a hydrogen bond with S46 of chain A, and Y262 forms a hydrogen bond with T116 of chain B. Additionally, D254 and Q275 form two salt bridges with R120 and E28, respectively, collectively stabilizing the scaffolding region (S8B Fig).

**Fig 4.**
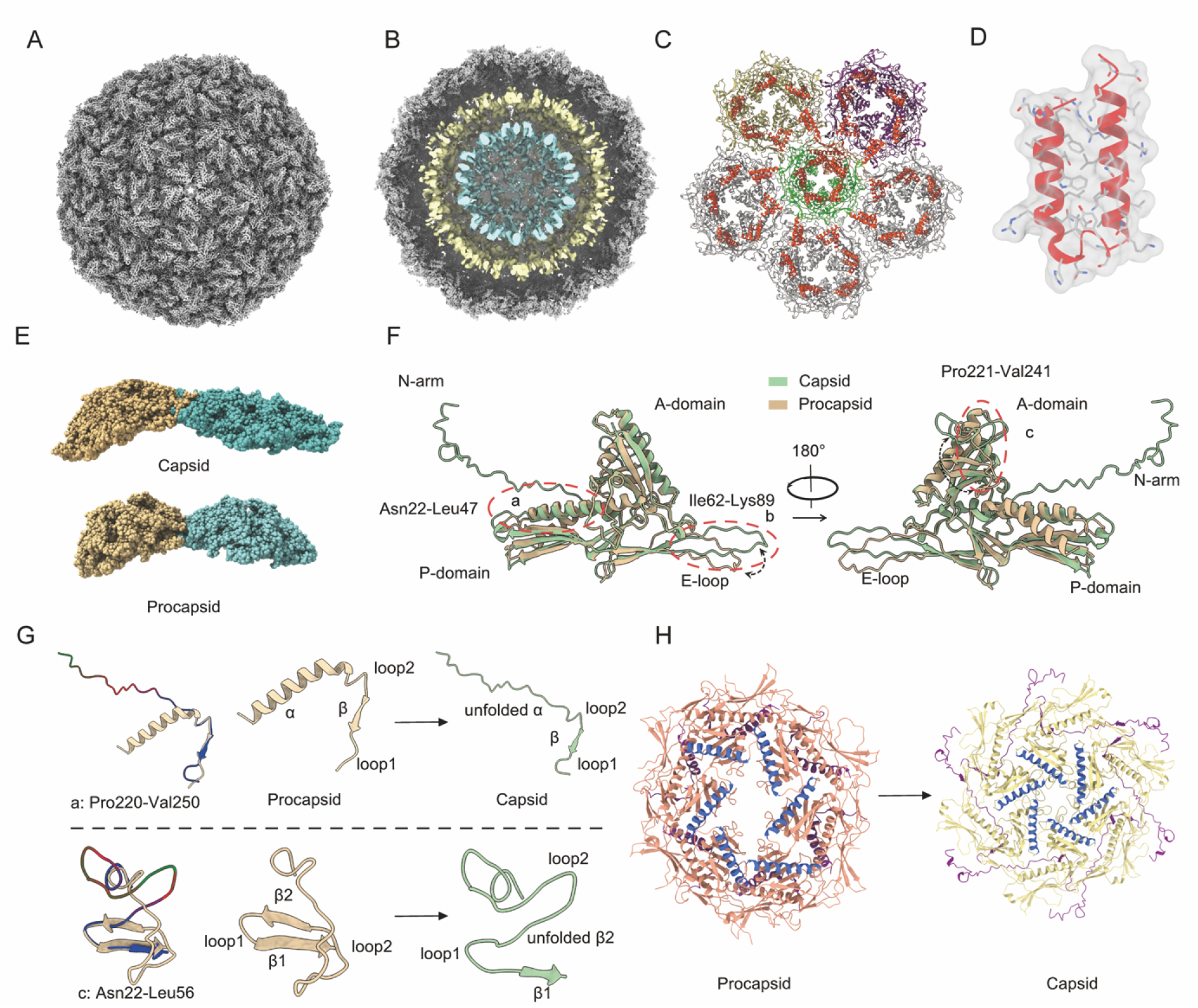
Cryo-EM structure of the procapsid and maturation-associated conformational transition. (**A**)Cryo-EM structure of the P22 procapsid. (**B**) Cryo-EM sectional view showing the double-layered concentric spherical structure inside the procapsid. The two layers are colored yellow and light blue, respectively. (**C**) Distribution of the U-shaped helix-turn-helix motif (C-terminus of the scaffolding protein) on the inner surface of the procapsid. (**D**) Atomic model of the C-terminal residues of the scaffolding protein fitted into the U-shaped helix-turn-helix density map. The density map is shown as a semi-transparent gray surface, and the atomic model is displayed as sticks, with the U-shaped helix-turn-helix motif highlighted in red. (**E**) Cross-sectional side views of the mature capsid and procapsid, showing expansion and elongation of the shell. (**F**) Superposition of the coat protein subunit from the procapsid and that from the mature capsid reveals the conformational changes that occur during capsid expansion. (**G**) Conformational transition of the N-arm and A-domain of the major capsid protein from the procapsid to the mature capsid. (**H**) Central aperture of the procapsid hexamer (left) and the mature capsid hexamer (right). Conformational changes at the A-domain tip are highlighted in blue, and those of the N-arm in purple.

In contrast to the mature capsid, the procapsid is less expanded, features an open six-fold aperture, and has a thicker shell (Fig 1B, Fig 4A and E). Conformational changes in gp37 trigger the structural rearrangements of the capsid. This is marked by alterations in the secondary structures of the P-domain, N-arm, and A-domain (Fig 4F), which exhibit significant RMSD fluctuations (S8C Fig). Key structural transitions include the unfolding of the N-arm α-helix (residues 220-250) that extends into the gp37 subunit of the neighboring asymmetric unit, and the unfolding of the A-domain β-sheet (residues 22-56) into an extended loop, which actively drags the extended α-helix to seal the six-fold axis aperture of the procapsid (Fig 4G-H).Taken together, we hypothesize that the conformational extension of the N-arm and A-domain drives the overall structural expansion of the capsid protein. These domain conformational changes promote the outward expansion of hexamers and pentamers (S1 Movie), ultimately forming the expanded mature capsid.

## Discussion

Despite the isolation of numerous *A. baumannii* phages and the disclosure of extensive genomic information in databases, structural information remains severely lacking. AbP20 is a podovirus that specifically infects multidrug-resistant *A. baumannii*. In the present work, we determined the high-resolution cryo-EM structure of AbP20 in its native state, providing detailed information that helps understand the assembly of the AbP20 capsid and the portal-tail complex. Our dissection of the entire AbP20 virion finely reveals the spatial organization of different protein components and deepens the understanding of the mechanisms that promote the stability and maturation of the phage capsid.

Capsid of tailed phages are primarily composed of the major capsid protein (MCP) and adopt a canonical HK97-like folding [11], including AbP20. Multiple copies of the major capsid protein form multimers (hexamers and pentamers) through the centrally clustered A-domain. Auxiliary domains, such as the P-domain, E-loop, and N-arm, facilitate the assembly of inter- and intra-capsid structures to form an icosahedral capsid due to their plasticity [28–30]. This structure is also retained in some eukaryotic DNA viruses [31]. Although sequence similarity is low, eukaryotic viruses have evolutionarily retained these classic and vital structural organization principles, expanding the diverse potential for the application and engineering of structural proteins.

Our reconstruction reveals the portal-tail gating system of AbP20. The portal and nozzle are responsible for DNA stabilization and injection, respectively. Although their specific charge properties are key characteristics that facilitate these interactions, the execution of their functions relies fundamentally on the overall structural context and spatial organization of the complex. The adaptor acts as an interaction hub among the tail fiber, portal, and nozzle, collaborating with other components to form a continuous, stable, and tightly controlled channel for DNA packaging and release. Although only the structure of the N-terminal residues 1-126 of the tail fiber is currently available, this region functions as its core anchoring module, firmly connecting the fiber to the tail complex and potentially maintaining its correct spatial orientation to enable the distal domains to perform subsequent functions.

Despite variations in capsid assembly strategies across different phages, the maturation and scaffold protein release mechanisms of AbP20 may much closely resemble those of P22 and T7. Previous biochemical studies have shown that scaffold protein gp8 of phage P22 plays an important role in the correct assembly of the procapsid [14]. Unlike P22 [17], the formation of AbP20 procapsid requires approximately a 1:1 ratio of capsid protein gp37 to scaffold protein gp36; however, this may not mean that every gp37 subunit is associated with one gp36 subunit but rather results from the double-layered concentric structure formed inside the procapsid. The expansion and maturation of the AbP20 capsid are likely triggered by a structural extension resulting from the unfolding of the gp37 N-arm α-helix and the A-domain β-strand. Conformational changes of the N-arm also occur in phages T4, T5, T7, SPP1, P22, P23-45, and λ [16, 17, 32-36]. Phages P22, SPP1, T7, T4, and λ encode scaffold proteins for capsid assembly. Among these phages, the scaffold proteins of P22, SPP1, and T7 are released through the hexamer apertures during capsid maturation [16, 18, 33]. In phages T4 and λ, the scaffold proteins are removed by virus-encoded proteases after capsid maturation [37, 38]. The mechanism of protease digestion in these phages explains why they lack a central aperture in the hexamer. Due to the similarities between AbP20 and T7/P22 in their assembly and capsid allosteric mechanisms, AbP20 may possess a similar scaffold protein release mechanism.

Although this study provides high-resolution structural insights into the *A. baumannii* phage AbP20, several limitations remain. Since the distal end of the tail fiber is dynamic and resolved at relatively low resolution, the detailed structure and function of the tail fiber remain unclear. Meanwhile, the gating mechanism during infection and the conformational changes among tail fibers still require extensive investigation. Although a common mechanism of capsid assembly and maturation is emerging, only two capsid forms have been resolved, and whether intermediate forms exist remains to be investigated. Further validation is needed to determine whether the portal participates in capsid packaging, interacts with the scaffolding protein, and contributes to the capsid maturation transition. In addition, the internal assembly and release of the scaffold protein, as well as what drives its release, still require in-depth exploration.

In summary, we have essentially elucidated the complete structure of AbP20, revealing the combinatorial organization of its five structural components. Structural analysis of the two capsid forms reveals a canonical pathway of capsid assembly and maturation. These exquisite structures greatly expand our understanding of the assembly of *A. baumannii* phage particles, particularly that of podovirus.

## Materials and Methods

### Phage purification

Phage AbP20 was added at an MOI of 0.01 to the exponential culture of *A. baumannii* strain Ab28 and cocultured for 4-6 h to allow phage proliferation. Afterwards, supernatant of the phage culture was harvested by centrifugation for 10 min at 8000 rpm, then 1 M NaCl and 10% PEG 8000 were added to the supernatant and stored at 4°C overnight for phage precipitation. Precipitated phage pellet collected by centrifugation at 10,000 × g for 1 h was resuspended in PBS and mixed with an equal volume of chloroform. Following centrifugation at 3500 rpm, phage was collected from the upper aqueous phase and carried out CsCl density gradient centrifugation to remove lipopolysaccharide and bacterial contaminants. Meanwhile, the upper band from the ultracentrifugation was collected to observe naturally occurring procapsids and mature capsids. The collected fractions were filtered through a 0.22 μm filter and stored at 4°C after extensive dialysis. Phage titer was determined by the plaque assay described below.

### Assembly and purification of the AbP20 procapsid

AbP20 genes encoding the scaffolding protein gp36 and the capsid protein gp37 were cloned into the pETDuet-1 expression vector (S1 Table) and transformed into *E. coli* BL21(DE3). Expression was induced overnight with isopropyl-beta-D-thiogalactopyranoside. Bacterial cells were harvested by centrifugation and resuspended in PBS. After sonication, cell debris was removed by centrifugation. NaCl and PEG 8000 were added to final concentrations of 1 M and 10%, respectively, followed by overnight precipitation at 4°C. The precipitated phage pellet was collected by centrifugation at 10,000 × g for 1 h, resuspended in PBS, and mixed with an equal volume of chloroform. Following centrifugation at 3,500 rpm, phages were collected from the upper aqueous phase and subjected to CsCl density gradient centrifugation to remove lipopolysaccharide and bacterial contaminants. The phage preparation was filtered through a 0.22 μm filter and stored at 4°C after extensive dialysis.

### Negative-stain transmission electron microscopy

A phage suspension at a concentration of 10^11^ PFU/ml was applied to a carbon-coated copper grid for 1 minute, followed by negative staining with 2.0% phosphotungstic acid (pH 7.0) for 2 minutes. Electron images were recorded on a Talos L120C transmission electron microscope.

### Cryo-EM sample preparation and data collection

To prepare the cryo-EM grids, 3 μL of purified AbP20 (10^12^ PFU/mL) and procapsid (1mg/mL) were applied to a freshly glow-discharged holey carbon grid (Quantifoil, BQR1.2/1.3-3C). After incubation for 10 seconds, the grid was blotted for 2 seconds at 4°C under 100% humidity before being plunge-frozen in liquid ethane using a Vitrobot Mark IV (Thermo Fisher Scientific). Cryo-EM single-particle data acquisition was performed on a CRYO ARM 300 transmission electron microscope (JEOL) operating at 300 kV with an in-column energy filter (slit width of 20 eV). Micrographs were collected automatically by a Gatan K3 direct electron detector in super-resolution mode using SerialEM [39], at a nominal magnification of 50,000× (corresponding to a pixel size of 0.475 Å). Each image stack was dose-fractionated to 40 frames with an accumulated dose of ~ 40 e/Å2 and a total exposure time of 1.6 s.

### Cryo-EM data processing

Image stacks were motion-corrected using MotionCor2 [40] and resulting dose-weighted. Micrographs were subjected to contrast transfer function (CTF) parameter estimation using CTFFIND4 [41]. A total of 101,289 AbP20 particles were automatically picked with Topaz [42] and subsequently extracted in RELION [43]. The particle extraction box size was 300 pixels, with each pixel corresponding to 3.80 Å. Following several rounds of two-dimensional (2D) and three-dimensional (3D) classification, 29,024 high-quality particles were re-extracted using a 608-pixel box size (1.25 Å per pixel). 3D refinement and postprocessing were performed in RELION with I3 icosahedral symmetry applied, yielding a 3.0-Å resolution density map of the AbP20 virion.

For reconstruction of the portal-tail complex, the 3.0-Å icosahedral reconstruction was first symmetry-expanded, followed by subparticle reconstruction of the twelve 5-fold vertices as described previously [44]. Vertex subparticles were then extracted with recentering offsets set to (0, 0, 300), using a 400-pixel box size (0.95 Å per pixel). After 3D classification, one class displaying clear tail structural features was selected for local refinement with C6 symmetry, resulting in a 3.3-Å resolution reconstruction. To obtain the complete tail-portal complex structure, re-extracted particles were centered at (0, 0, 200) with a 400-pixel box size (1.9 Å per pixel). The final tail map was reconstructed at a resolution of 4.4 Å. Following a similar processing procedure to that of AbP20, 43,377 self-assembled capsid particles were selected for 3D reconstruction with I3 symmetry at a resolution of 3.3 Å. All structure resolutions were estimated based on the gold standard Fourier Shell Correlation (FSC) = 0.143 criterion.

### Atomic model building and model refinement

Initial atomic models of the capsid protein (gp37), scaffolding protein (gp36), and tail-portal complex components (portal protein gp35, adaptor protein gp39, nozzle protein gp40, and fiber protein gp44) were generated using AlphaFold2 [45]. All models were docked into the density maps in UCSF Chimera [46] and subsequently manually adjusted in Coot [47]. The refined models were then subjected to real-space refinement with Phenix [48]. The asymmetric unit of the AbP20 capsid or self-assembled capsid comprises seven copies of capsid protein. Models of the adaptor, N-terminal region of the fiber, nozzle, and portal were refined based on the density map of the AbP20 portal-tail complex. The C-terminal atomic structure of the fiber protein was solved by X-ray crystallography. All structural figures were prepared using ChimeraX. Detailed information on cryo-EM data collection, processing, and refinement is summarized in S2 Table.

### Structural comparison

Published capsid amino acid sequences were obtained from UniProt and aligned using ClustalW [49]. Asymmetric unit structural models were downloaded from the PDB website and structurally compared using ChimeraX.

## Acknowledgments

We are grateful to Ding Gao and Bi-Chao Xu from the Core Facility and Technical Support, Wuhan Institute of Virology for their assistance in image and transmission electron microscopy.

## Data availability

Coordinates and structure factors of the capsid and the tail structure of phage AbP20, and the assembled structure of the capsid, have been deposited in PDB under the accession codes 9XMS, 9XMU, and 9XMT, respectively.

## List of legends for Supporting Information

**Fig. S1. Morphological observation of AbP20 virions**. (**A**) Representative negative-stain micrograph of AbP20 virions. Scale bar, 200 nm. (**B**) Representative cryo-EM micrograph of AbP20 virions. Scale bar, 100 nm.

**Fig. S2. Cryo-EM data processing of the AbP20 virion**. Subparticle reconstructions of the capsid and portal-tail complex are presented as a flowchart, indicating the resolutions and numbers of phage particles used.

**Fig. S3. Comparison of major capsid proteins**. (**A**) Sequence alignment of major capsid proteins. (**B**) Structural comparison of the AbP20 major capsid protein with those of HK97 (PDB: 1OHG), T7 (PDB: 3J7W), and HSV (residues 1-398; PDB: 6CGR).

**Fig. S4. The subunit interactions within/between capsomeres of the AbP20 capsid**. (**A**) The overall view of one penton (green) and two hexons (pale pink) of the AbP20 capsid. (**B-D**) The interaction details shown in the dashed boxes are displayed at higher magnification in the following panels, including the interactions between the penton and two hexons (B), between pentons (C), and between hexons (D). Different domains of the major capsid protein are colored.

**Fig. S5. The structure of AbP20 portal-tail complex**. (**A**) Central channel of the gating complex. (**B**) Electrostatic potential of the central channel of the gating complex (left); side and top views of the electrostatic potential of the gating complex (right). Negatively and positively charged regions are colored red and blue, respectively. (**C**) Monomeric structures of the gating components. Different domains are colored differently.

**Fig. S6. Morphological observation of AbP20 procapsid**. (**A**) Representative cryo-electron micrograph of self-assembled AbP20 procapsids. Scale bar, 100 nm. (B) Cryo-EM images of naturally isolated AbP20 procapsids and mature capsids. Scale bar, 100 nm. The red arrows represent procapsid, and the yellow arrows represent mature capsid.

**Fig. S7. Cryo-EM data processing of the AbP20 procapsid**. Subparticle reconstruction of the procapsid is presented as a flowchart, indicating the resolutions and numbers of phage particles used.

**Fig. S8. Capsid maturation and interactions between the scaffolding protein and the procapsid**. (**A**) Cryo-EM density map showing the double-layered concentric spherical structure inside the procapsid. The two layers are colored yellow and light blue, respectively.(**B**)Interactions between the C-terminus of the scaffolding protein and the major capsid protein. High-magnification views show the interaction details of the five interfaces, which are highlighted with dashed boxes. Residues involved in the interactions are shown as sticks and labeled. Hydrogen bonds and salt bridges are indicated by blue and red dashed lines, respectively. (**C**) Structural superimposition of the capsid protein subunits from the procapsid and mature capsid. The mature capsid is color-mapped based on RMSD values (blue to green gradient). The A-domain, N-arm, E-loop, and P-domain are indicated.

**Table S1. Primers used in this study**.

**Table S2. Cryo-EM data collection, processing, and model refinement**

**PDB Validation reports for the capsid and the tail structure of phage AbP20, and the assembled structure of the capsid under the accession codes 9XMS, 9XMU, and 9XMT**.

## References

1. Moubareck CA, Halat DH. Insights into : A Review of Microbiological, Virulence, and Resistance Traits in a Threatening Nosocomial Pathogen. Antibiotics-Basel. 2020;9(3). doi: ARTN 11910.3390/antibiotics9030119. PubMed PMID: WOS:000523481200002.

2. Wu N, Dai J, Guo M, Li J, Zhou X, Li F, et al. Pre-optimized phage therapy on secondary Acinetobacter baumannii infection in four critical COVID-19 patients. Emerg Microbes Infect. 2021;10(1):612–8. doi: 10.1080/22221751.2021.1902754. PubMed PMID: 33703996; PubMed Central PMCID: PMCPMC8032346.

3. LaVergne S, Hamilton T, Biswas B, Kumaraswamy M, Schooley RT, Wooten D. Phage Therapy for a Multidrug-Resistant Acinetobacter baumannii Craniectomy Site Infection. Open Forum Infect Dis. 2018;5(4):ofy064. Epub 20180323. doi: 10.1093/ofid/ofy064. PubMed PMID: 29687015; PubMed Central PMCID: PMCPMC5905571.

4. Rao S, Betancourt-Garcia M, Kare-Opaneye YO, Swierczewski BE, Bennett JW, Horne B, et al. Critically Ill Patient with Multidrug-Resistant Acinetobacter baumannii Respiratory Infection Successfully Treated with Intravenous and Nebulized Bacteriophage Therapy. Antimicrob Agents Chemother. 2022;66(1):e0082421. Epub 20211018. doi: 10.1128/AAC.00824-21. PubMed PMID: 34662188; PubMed Central PMCID: PMCPMC8765271.

5. Eales BM, Tam VH. Case Commentary: Novel Therapy for Multidrug-Resistant Acinetobacter baumannii Infection. Antimicrob Agents Chemother. 2022;66(1):e0199621. Epub 20211025. doi: 10.1128/AAC.01996-21. PubMed PMID: 34694874; PubMed Central PMCID: PMCPMC8765303.

6. Meng R, Xing Z, Chang JY, Yu Z, Thongchol J, Xiao W, et al. Structural basis of Acinetobacter type IV pili targeting by an RNA virus. Nat Commun. 2024;15(1):2746. Epub 20240329. doi: 10.1038/s41467-024-47119-5. PubMed PMID: 38553443; PubMed Central PMCID: PMCPMC10980823.

7. Alseth EO, Roush C, Irby I, Kopylov M, Bobe D, Diggs MW, et al. Mystique, a broad host range Acinetobacter phage, reveals the impact of culturing conditions on phage isolation and infectivity. PLoS Pathog. 2025;21(4):e1012986. Epub 20250410. doi: 10.1371/journal.ppat.1012986. PubMed PMID: 40208916; PubMed Central PMCID: PMCPMC12013898.

8. Steven AC, Heymann JB, Cheng NQ, Trus BL, Conway JF. Virus maturation: dynamics and mechanism of a stabilizing structural transition that leads to infectivity. Curr Opin Struc Biol. 2005;15(2):227–36. doi: 10.1016/j.sbi.2005.03.008. PubMed PMID: WOS:000229065800015.

9. Aksyuk AA, Rossmann MG. Bacteriophage assembly. Viruses. 2011;3(3):172–203. Epub 20110225. doi: 10.3390/v3030172. PubMed PMID: 21994726; PubMed Central PMCID: PMCPMC3185693.

10. Droge A, Santos MA, Stiege AC, Alonso JC, Lurz R, Trautner TA, et al. Shape and DNA packaging activity of bacteriophage SPP1 procapsid: protein components and interactions during assembly. J Mol Biol. 2000;296(1):117–32. doi: 10.1006/jmbi.1999.3450. PubMed PMID: 10656821.

11. Oh B, Moyer CL, Hendrix RW, Duda RL. The delta domain of the HK97 major capsid protein is essential for assembly. Virology. 2014;456-457:171–8. Epub 20140410. doi: 10.1016/j.virol.2014.03.022. PubMed PMID: 24889236; PubMed Central PMCID: PMCPMC4044616.

12. Huet A, Conway JF, Letellier L, Boulanger P. In vitro assembly of the T=13 procapsid of bacteriophage T5 with its scaffolding domain. J Virol. 2010;84(18):9350–8. Epub 20100623. doi: 10.1128/JVI.00942-10. PubMed PMID: 20573812; PubMed Central PMCID: PMCPMC2937613.

13. Choi KH, Morais MC, Anderson DL, Rossmann MG. Determinants of bacteriophage phi29 head morphology. Structure. 2006;14(11):1723–7. doi: 10.1016/j.str.2006.09.007. PubMed PMID: 17098197.

14. Earnshaw W, King J. Structure of phage P22 coat protein aggregates formed in the absence of the scaffolding protein. J Mol Biol. 1978;126(4):721–47. doi: 10.1016/0022-2836(78)90017-7. PubMed PMID: 370407.

15. Ray P, Murialdo H. The role of gene Nu3 in bacteriophage lambda head morphogenesis. Virology. 1975;64(1):247–63. doi: 10.1016/0042-6822(75)90096-3. PubMed PMID: 1090075.

16. Guo F, Liu Z, Fang PA, Zhang Q, Wright ET, Wu W, et al. Capsid expansion mechanism of bacteriophage T7 revealed by multistate atomic models derived from cryo-EM reconstructions. Proc Natl Acad Sci U S A. 2014;111(43):E4606–14. Epub 20141013. doi: 10.1073/pnas.1407020111. PubMed PMID: 25313071; PubMed Central PMCID: PMCPMC4217468.

17. Xiao H, Zhou J, Yang F, Liu Z, Song J, Chen W, et al. Assembly and Capsid Expansion Mechanism of Bacteriophage P22 Revealed by High-Resolution Cryo-EM Structures. Viruses. 2023;15(2). Epub 20230126. doi: 10.3390/v15020355. PubMed PMID: 36851569; PubMed Central PMCID: PMCPMC9965877.

18. Chen DH, Baker ML, Hryc CF, DiMaio F, Jakana J, Wu W, et al. Structural basis for scaffolding-mediated assembly and maturation of a dsDNA virus. Proc Natl Acad Sci U S A. 2011;108(4):1355–60. Epub 20110110. doi: 10.1073/pnas.1015739108. PubMed PMID: 21220301; PubMed Central PMCID: PMCPMC3029737.

19. Xue H, Li X, Rao G, Hu F, Zhong M, Miernikiewicz P, et al. Antibodies targeting phage fiber and nozzle proteins impair Acinetobacter baumannii phage therapy by blocking infection and promoting immune clearance. PLoS Biol. 2026;24(9):e3004009. Epub 2026/09/22 21:36. doi: 10.1371/journal.pbio.3004009. PubMed PMID: 42771684; PubMed Central PMCID: PMCPMC13623143.

20. Liu XA, Zhang QF, Murata K, Baker ML, Sullivan MB, Fu C, et al. Structural changes in a marine podovirus associated with release of its genome into. Nat Struct Mol Biol. 2010;17(7):830–U76. doi: 10.1038/nsmb.1823. PubMed PMID: WOS:000279631500009.

21. Chen WY, Xiao H, Wang XR, Song SL, Han Z, Li XW, et al. Structural changes of a bacteriophage upon DNA packaging and maturation. Protein Cell. 2020;11(5):374–9. doi: 10.1007/s13238-020-00715-9. PubMed PMID: WOS:000524402700001.

22. Huet A, Oh B, Maurer J, Duda RL, Conway JF. A symmetry mismatch unraveled: How phage HK97 scaffold flexibly accommodates a 12-fold pore at a 5-fold viral capsid vertex. Sci Adv. 2023;9(24):eadg8868. Epub 20230616. doi: 10.1126/sciadv.adg8868. PubMed PMID: 37327331; PubMed Central PMCID: PMCPMC10275583.

23. Wikoff WR, Liljas L, Duda RL, Tsuruta H, Hendrix RW, Johnson JE. Topologically linked protein rings in the bacteriophage HK97 capsid. Science. 2000;289(5487):2129–33. doi: 10.1126/science.289.5487.2129. PubMed PMID: 11000116.

24. Dai X, Zhou ZH. Structure of the herpes simplex virus 1 capsid with associated tegument protein complexes. Science. 2018;360(6384). Epub 20180405. doi: 10.1126/science.aao7298. PubMed PMID: 29622628; PubMed Central PMCID: PMCPMC5959011.

25. Dedeo CL, Cingolani G, Teschke CM. Portal Protein: The Orchestrator of Capsid Assembly for the dsDNA Tailed Bacteriophages and Herpesviruses. Annu Rev Virol. 2019;6(1):141–60. Epub 20190723. doi: 10.1146/annurev-virology-092818-015819. PubMed PMID: 31337287; PubMed Central PMCID: PMCPMC6947915.

26. Cuervo A, Pulido-Cid M, Chagoyen M, Arranz R, Gonzalez-Garcia VA, Garcia-Doval C, et al. Structural characterization of the bacteriophage T7 tail machinery. J Biol Chem. 2013;288(36):26290–9. Epub 20130724. doi: 10.1074/jbc.M113.491209. PubMed PMID: 23884409; PubMed Central PMCID: PMCPMC3764833.

27. Cuervo A, Fabrega-Ferrer M, Machon C, Conesa JJ, Fernandez FJ, Perez-Luque R, et al. Structures of T7 bacteriophage portal and tail suggest a viral DNA retention and ejection mechanism. Nat Commun. 2019;10(1):3746. Epub 20190820. doi: 10.1038/s41467-019-11705-9. PubMed PMID: 31431626; PubMed Central PMCID: PMCPMC6702177.

28. Helgstrand C, Wikoff WR, Duda RL, Hendrix RW, Johnson JE, Liljas L. The refined structure of a protein catenane: the HK97 bacteriophage capsid at 3.44 A resolution. J Mol Biol. 2003;334(5):885–99. doi: 10.1016/j.jmb.2003.09.035. PubMed PMID: 14643655.

29. Stone NP, Demo G, Agnello E, Kelch BA. Principles for enhancing virus capsid capacity and stability from a thermophilic virus capsid structure. Nat Commun. 2019;10(1):4471. Epub 20191002. doi: 10.1038/s41467-019-12341-z. PubMed PMID: 31578335; PubMed Central PMCID: PMCPMC6775164.

30. Cai L, Liu H, Zhang W, Xiao S, Zeng Q, Dang S. Cryo-EM structure of cyanophage P-SCSP1u offers insights into DNA gating and evolution of T7-like viruses. Nat Commun. 2023;14(1):6438. Epub 20231013. doi: 10.1038/s41467-023-42258-7. PubMed PMID: 37833330; PubMed Central PMCID: PMCPMC10575957.

31. Newcomb WW, Thomsen DR, Homa FL, Brown JC. Assembly of the herpes simplex virus capsid: Identification of soluble scaffold-portal complexes and their role in formation of portal-containing capsids. Journal of Virology. 2003;77(18):9862–71. doi: 10.1128/Jvi.77.18.9862-9871.2003. PubMed PMID: WOS:000185046200015.

32. Zinke M, Schroder GF, Lange A. Major tail proteins of bacteriophages of the order Caudovirales. J Biol Chem. 2022;298(1):101472. Epub 20211208. doi: 10.1016/j.jbc.2021.101472. PubMed PMID: 34890646; PubMed Central PMCID: PMCPMC8718954.

33. Ignatiou A, Brasiles S, El Sadek Fadel M, Burger J, Mielke T, Topf M, et al. Structural transitions during the scaffolding-driven assembly of a viral capsid. Nat Commun. 2019;10(1):4840. Epub 20191024. doi: 10.1038/s41467-019-12790-6. PubMed PMID: 31649265; PubMed Central PMCID: PMCPMC6813328.

34. Bayfield OW, Klimuk E, Winkler DC, Hesketh EL, Chechik M, Cheng N, et al. Cryo-EM structure and in vitro DNA packaging of a thermophilic virus with supersized T=7 capsids. Proc Natl Acad Sci U S A. 2019;116(9):3556–61. Epub 20190208. doi: 10.1073/pnas.1813204116. PubMed PMID: 30737287; PubMed Central PMCID: PMCPMC6397560.

35. Huet A, Duda RL, Boulanger P, Conway JF. Capsid expansion of bacteriophage T5 revealed by high resolution cryoelectron microscopy. Proc Natl Acad Sci U S A. 2019;116(42):21037–46. Epub 20191002. doi: 10.1073/pnas.1909645116. PubMed PMID: 31578255; PubMed Central PMCID: PMCPMC6800373.

36. Fang Q, Tang WC, Fokine A, Mahalingam M, Shao Q, Rossmann MG, et al. Structures of a large prolate virus capsid in unexpanded and expanded states generate insights into the icosahedral virus assembly. Proc Natl Acad Sci U S A. 2022;119(40):e2203272119. Epub 20220926. doi: 10.1073/pnas.2203272119. PubMed PMID: 36161892; PubMed Central PMCID: PMCPMC9546572.

37. Medina E, Wieczorek D, Medina EM, Yang Q, Feiss M, Catalano CE. Assembly and maturation of the bacteriophage lambda procapsid: gpC is the viral protease. J Mol Biol. 2010;401(5):813–30. Epub 20100708. doi: 10.1016/j.jmb.2010.06.060. PubMed PMID: 20620152.

38. Fokine A, Rossmann MG. Common Evolutionary Origin of Procapsid Proteases, Phage Tail Tubes, and Tubes of Bacterial Type VI Secretion Systems. Structure. 2016;24(11):1928–35. Epub 20160922. doi: 10.1016/j.str.2016.08.013. PubMed PMID: 27667692; PubMed Central PMCID: PMCPMC5093050.

39. de la Cruz MJ, Martynowycz MW, Hattne J, Gonen T. MicroED data collection with SerialEM. Ultramicroscopy. 2019;201:77–80. Epub 20190319. doi: 10.1016/j.ultramic.2019.03.009. PubMed PMID: 30986656; PubMed Central PMCID: PMCPMC6752703.

40. Zheng SQ, Palovcak E, Armache JP, Verba KA, Cheng Y, Agard DA. MotionCor2: anisotropic correction of beam-induced motion for improved cryo-electron microscopy. Nat Methods. 2017;14(4):331–2. Epub 20170227. doi: 10.1038/nmeth.4193. PubMed PMID: 28250466; PubMed Central PMCID: PMCPMC5494038.

41. Rohou A, Grigorieff N. CTFFIND4: Fast and accurate defocus estimation from electron micrographs. J Struct Biol. 2015;192(2):216–21. Epub 20150813. doi: 10.1016/j.jsb.2015.08.008. PubMed PMID: 26278980; PubMed Central PMCID: PMCPMC6760662.

42. Bepler T, Morin A, Rapp M, Brasch J, Shapiro L, Noble AJ, et al. Positive-unlabeled convolutional neural networks for particle picking in cryo-electron micrographs. Nat Methods. 2019;16(11):1153–60. Epub 20191007. doi: 10.1038/s41592-019-0575-8. PubMed PMID: 31591578; PubMed Central PMCID: PMCPMC6858545.

43. Scheres SH. RELION: implementation of a Bayesian approach to cryo-EM structure determination. J Struct Biol. 2012;180(3):519–30. Epub 20120919. doi: 10.1016/j.jsb.2012.09.006. PubMed PMID: 23000701; PubMed Central PMCID: PMCPMC3690530.

44. Pan M, Alvarez-Cabrera AL, Kang JS, Wang L, Fan C, Zhou ZH. Asymmetric reconstruction of mammalian reovirus reveals interactions among RNA, transcriptional factor micro2 and capsid proteins. Nat Commun. 2021;12(1):4176. Epub 20210707. doi: 10.1038/s41467-021-24455-4. PubMed PMID: 34234134; PubMed Central PMCID: PMCPMC8263624.

45. Jumper J, Evans R, Pritzel A, Green T, Figurnov M, Ronneberger O, et al. Highly accurate protein structure prediction with AlphaFold. Nature. 2021;596(7873):583–9. Epub 20210715. doi: 10.1038/s41586-021-03819-2. PubMed PMID: 34265844; PubMed Central PMCID: PMCPMC8371605.

46. Pettersen EF, Goddard TD, Huang CC, Meng EC, Couch GS, Croll TI, et al. UCSF ChimeraX: Structure visualization for researchers, educators, and developers. Protein Sci. 2021;30(1):70–82. Epub 20201022. doi: 10.1002/pro.3943. PubMed PMID: 32881101; PubMed Central PMCID: PMCPMC7737788.

47. Emsley P, Cowtan K. Coot: model-building tools for molecular graphics. Acta Crystallogr D Biol Crystallogr. 2004;60(Pt 12 Pt 1):2126–32. Epub 20041126. doi: 10.1107/S0907444904019158. PubMed PMID: 15572765.

48. Adams PD, Afonine PV, Bunkoczi G, Chen VB, Davis IW, Echols N, et al. PHENIX: a comprehensive Python-based system for macromolecular structure solution. Acta Crystallogr D Biol Crystallogr. 2010;66(Pt 2):213–21. Epub 20100122. doi: 10.1107/S0907444909052925. PubMed PMID: 20124702; PubMed Central PMCID: PMCPMC2815670.

49. Thompson JD, Gibson TJ, Higgins DG. Multiple sequence alignment using ClustalW and ClustalX. Curr Protoc Bioinformatics. 2002;Chapter 2:Unit 2 3. doi: 10.1002/0471250953.bi0203s00. PubMed PMID: 18792934.

